# Domain Insertion Improves the Precision of a CRISPR Adenine Base Editor

**DOI:** 10.64898/2026.07.03.736350

**Authors:** Marik M. Müller, Nicholas T. Southern, Dominik Niopek

## Abstract

Adenine base editors (ABEs) enable efficient A•T to G•C conversion, but their broad activity windows frequently cause unintended bystander edits. We hypothesized that insertion of a bulky, inert protein domain into the base editor would limit the effective reach of the deaminase, thereby preferentially directing editing to the intended target adenine.

Here, we systematically map domain insertion sites within the high-activity TadA8e adenine base editor using structure-guided design and computational inference. We find that TadA8e accepts domain insertions at multiple surface sites, with overall activity and editing window width dependent on insert position rather than domain identity. Excitingly, inserting domains at residue L68 preserved robust editing across multiple genomic loci while tightly focusing the editing window around position 5. Insertion of superfolder GFP at this site produced a base editor variant with a narrower editing window, the ability to track edited cells by fluorescence, and markedly reduced Cas-independent off-target editing. Our work highlights domain insertion engineering as a powerful strategy to create more focused and precise CRISPR base editors.

## Introduction

Adenine base editors (ABEs) enable programmable A•T to G•C conversions in genomic DNA without introducing double-strand breaks and have become central tools for precise genome engineering^1^. These editor complexes typically consist of an *Escherichia coli*-derived TadA deaminase domain fused to a Cas nickase, most commonly Cas9 from *Streptococcus pyogenes*^*2,3*^ (Figure 1A). Iterative evolution and rational engineering of the TadA domain have produced a suite of highly active deaminases^2–6^. Among these, the extensively evolved TadA8e variant exhibits high editing efficiencies across a wide range of cellular contexts^4^. However, this increased catalytic activity is accompanied by broadened editing windows that typically span multiple nucleotides within the protospacer, frequently resulting in unintended bystander mutations^4,7^. Beyond its broad editing window, the high catalytic activity and increased processivity of TadA8e renders ABE8e more susceptible to Cas-dependent as well as Cas-independent off-target editing, including transcriptome-wide RNA editing and deamination of exposed ssDNA substrates^4^.

**Figure 1.**
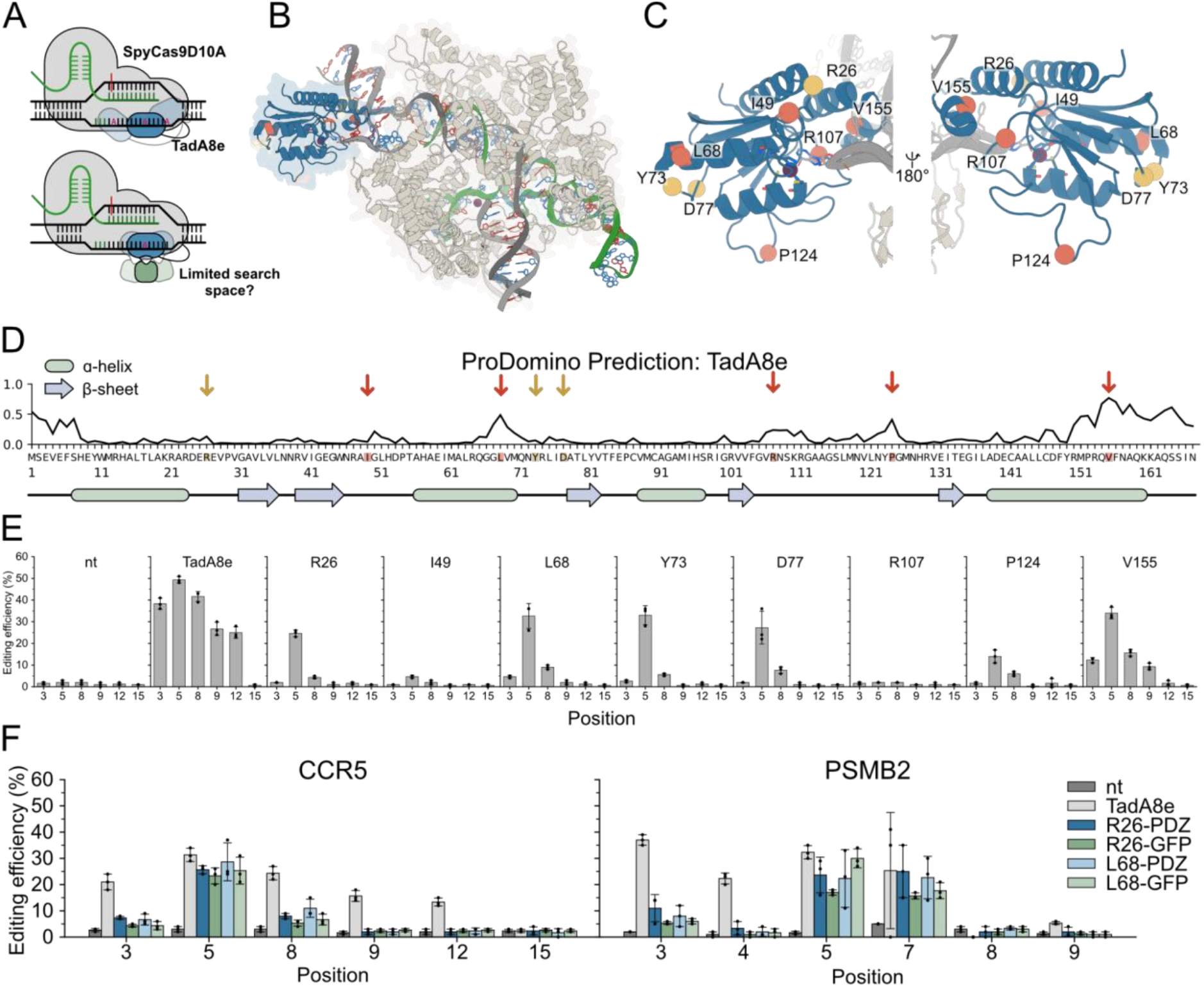
TadA8e tolerates domain insertion at multiple sites. (A) Working model for editing window narrowing. In the unmodified editor (top), the deaminase samples a broad stretch of the non-target ssDNA exposed within the R-loop. Insertion of a bulky domain (bottom) restricts this sampling, focusing editing on the target adenine. (B) Structure of ABE8e bound to DNA (PDB 6VPC): The TadA8e deaminase is in dark blue, the *Spy*Cas9 nickase in light blue, the gRNA in green, the target strand in dark grey, and the non-target strand in light grey color. Insertion sites are colored orange (ProDomino-predicted) or yellow (manually selected). (C) Close-up of the TadA8e domain shown in two orientations, insertion sites are labeled and colored as in (B). (D) ProDomino trace of TadA8e depicting the predicted insertion tolerance for each residue. Highlighted sites were chosen for testing. Secondary structure elements, α-helices (green) and β-sheets (blue), are indicated below. (E) Bar charts showing A→G editing at the endogenous CCR5 locus in HEK293T cells. Cells expressing either ABE8e-*As*LOV2 hybrids, with the LOV2 inserted at the indicated sites in TadA8e, or wild-type ABE8e as well as a CCR5-targeting sgRNA were incubated for 48 h, followed by sanger sequencing and EditR analysis. (F) A→G Editing at CCR5 and PSMB2 with PDZ or sfGFP inserted after position R26 or L68 within TadA8e of ABE8e. Numbers indicate the position of each target adenine within the protospacer, counted from the PAM-distal end (position 1). (E, F) Data are mean ± SD of n = 3 independent experiments.

Extensive engineering efforts have sought to narrow the editing window and improve positional precision, most commonly by tuning the deaminase domain through point mutations^6,8^. These approaches have been successful and can substantially reduce bystander editing. However, improved precision achieved through weakening or altering deaminase activity may come at the cost of reduced overall editing efficiency^6,8^. Moreover, point mutations do not necessarily address a key physical contributor to bystander editing: the ability of TadA to spatially sample multiple adenines within a longer single-stranded DNA region exposed upon R-loop formation. We hypothesized that inserting larger, autonomously folding domains into the editor architecture could constrain the spatial reach of the deaminase by mechanically tuning TadA8e mobility, thereby narrowing the editing window while preserving high editing activity at spatially preferred target sites.

Here, using computational inference and structure-based determination of permissive insertion sites^9^, we investigate TadA8e’s tolerance to larger domain inserts and assess its effects on the editing window width as well as on Cas-independent off-target editing.

## Results

### TadA8e Tolerates Domain Insertion Across Distinct Surface Sites

To systematically map the tolerance of the TadA8e deaminase to internal domain insertion, a combinatorial approach utilizing structure-guided design and computational predictions was used. We selected three domain insertion sites within flexible, surface-exposed loops (R26, Y73, D77) and used the machine learning algorithm ProDomino to predict five additional sites, located primarily in alpha-helices (I49, L68, R107, P124, V155) (Figure 1B, C, D). Each site received an insertion of the *Avena sativa* light-oxygen-voltage domain (LOV2), chosen because its closely spaced N- and C-termini minimize structural distortion upon insertion and because blue light induced unfolding of its Jα helix offered the possibility of assessing optogenetic modulation of base editor activity in parallel^10,11^.

We co-expressed the resulting Cas9-TadA8e-LOV2 hybrids with a *CCR5*-targeting sgRNA in HEK293T cells via transient transfection and assessed the overall editing efficiency as well as the editing window (Figure 1E). Six out of the eight tested insertion sites accommodated the *As*LOV2 domain well overall and retained substantial catalytic activity, with the best performing variant (TadA8e-L68-LOV2) showing an editing efficiency as high as ∼80 % or more of the wild-type TadA8e, dependent on the experiment. Of note, none of the hybrids conferred strong light-dependent changes in both editing efficacy and window width (Supplementary Figure S2), suggesting a lack of allosteric wiring between the chosen insertion sites and the catalytic surface of TadA8e. Jointly, these results demonstrate that the TadA8e scaffold is broadly permissive to domain insertion, but does not facilitate allosteric coupling - at least when using LOV2 as sensory domain.

### Domain Insertion Narrows the Editing Window

Beyond domain insertion tolerance, our initial screen revealed that the LOV2 domain insertion strongly altered the positional editing profile of TadA8e as hypothesized. Hybrids generated through domain insertion at R26, L68, Y73, and D77 of TadA8e preserved high editing efficiency at the central target position A5 while strongly reducing activity at all bystander adenines within the editing window (Figure 1E). The V155 variant matched the editing efficiency of these hybrids at A5 but achieved only modest suppression of bystander activity.

To determine whether the narrowed editing window reflected a specific property of LOV2 or a general consequence of domain insertion into TadA8e, we exchanged LOV2 in two of our most promising hybrids (R26 and L68) with structurally distinct globular domains: a functionally inert PDZ domain and the larger, fluorescent superfolder GFP (sfGFP). All four domain insertions were well-tolerated and resulted in similarly narrowed editing windows to those of initial LOV2 variants for CCR5, a trend that was recapitulated at a second, independent locus (PSMB2) (Figure 1F). Remarkably, the L68-sfGFP variant retained near wild-type activity at A5 (CCR5) and A5/A7 (PSMB2) while strongly reducing editing at bystander sites. Together, these results suggest that the observed narrowing of the editing window is not simply a consequence of reduced base-editor efficiency, but rather reflects structural constraints imposed by the inserted domain that limit deaminase access to the exposed DNA substrate.

### Insertion of sfGFP Yields a Fluorescent Editor with a Narrowed Editing Window and Increased Locus Specificity

Beyond its effect on editing specificity, the L68-sfGFP variant offers a practical advantage: integrating sfGFP directly into the deaminase allows for fluorescence-based tracking while likewise reducing bystander editing. Indeed, fluorescence microscopy confirmed that the internal sfGFP domain folds correctly within the TadA8e-L68 scaffold, producing a fluorescent signal that localized to the nucleus in living cells (Figure 2A).

**Figure 2.**
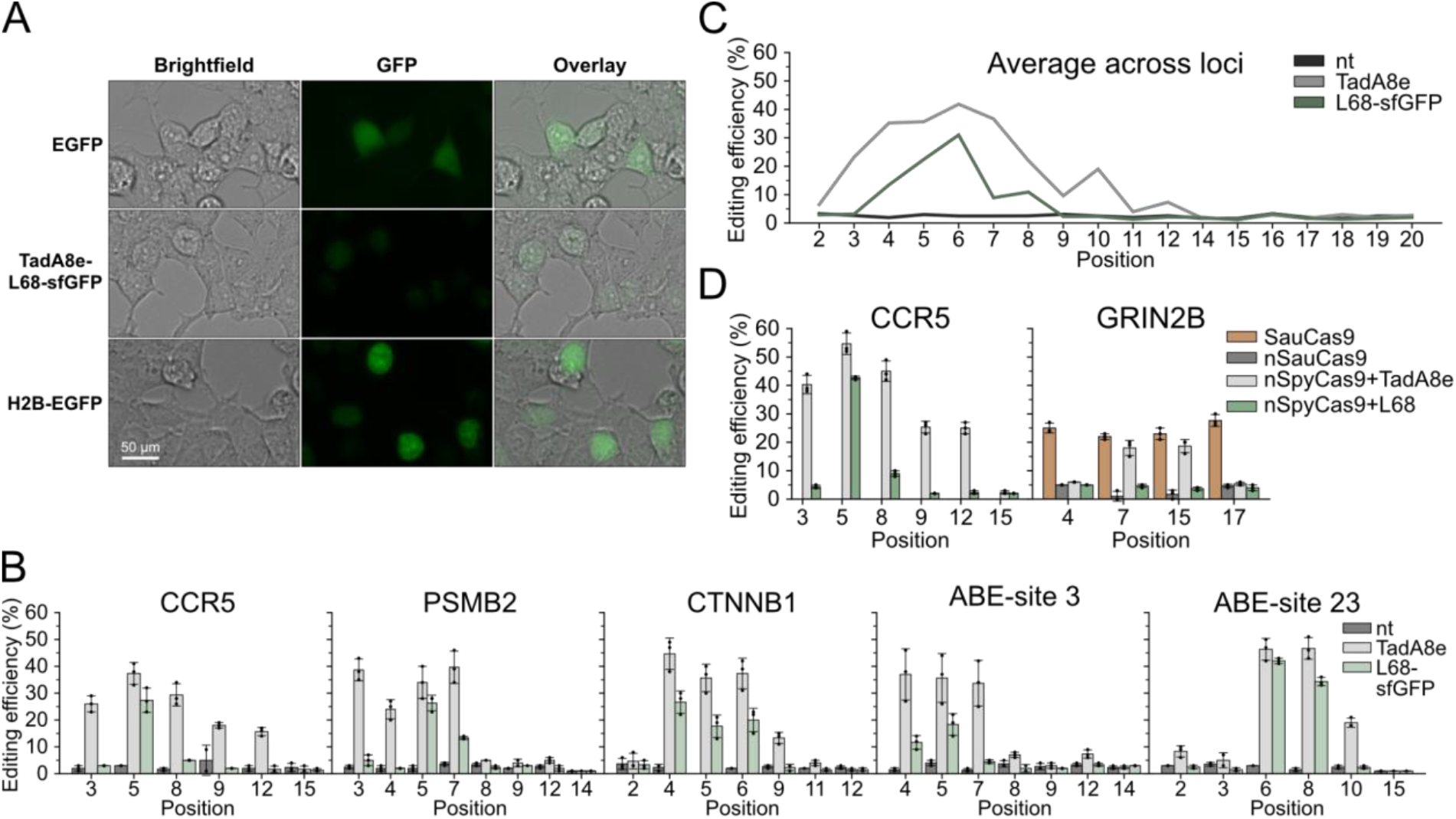
TadA8e-L68-sfGFP is a fluorescent base editor with a constricted editing window and low Cas-independent off-target activity. (A) Live-cell fluorescence microscopy of HEK293T cells expressing EGFP (top), TadA8e-L68-sfGFP (middle), or H2B-EGFP (bottom); columns show brightfield, GFP, and overlay. GFP exposure: 0.2 s (TadA8e-L68-sfGFP, H2B-EGFP), 0.5 s (EGFP control). Scale bar, 50 µm. (B) Per-position A→G editing at five endogenous loci (*CCR5, PSMB2*, ABE-site 3, *CTNNB1*, ABE-site 23) in HEK293T cells, comparing wild-type TadA8e and TadA8e-L68-sfGFP against a non-targeting control (nt). Editing efficiencies were quantified from Sanger sequencing traces using EditR. Mean ± SD of n = 3 independent experiments. (C) A→G editing averaged across the five loci taken from (B) to estimate editing window size. (D) Orthogonal R-loop assay to assess Cas-independent off-target deamination. The nSpyCas9 adenine base editor, containing either wild-type TadA8e or the L68-sfGFP insertion variant, was programmed to edit the CCR5 on-target locus. In parallel, an n*Sau*Cas9 nickase was targeted to GRIN2B to generate an orthogonal R-loop and expose single-stranded DNA at a potential trans off-target site. A *Sau*Cas9 adenine base editor targeted to GRIN2B served as a control for editing competence at this locus. Numbers denote the position of each target adenine within the protospacer, counted from the PAM-distal end, with the first nucleotide defined as position 1. Mean ± SD of n = 3 independent experiments.

To comprehensively profile the editing behavior of the TadA8e-L68-sfGFP variant, we assessed its activity across five distinct genomic loci. The observed editing efficacy was comparable to wild-type TadA8e at the key position A5/A6 (counting the pam distal site of the spacer as 1) for three out of the five tested sites (Figure 2B). Analyzing the averaged editing activity across all loci revealed that wild-type TadA8e possesses a broad editing window spanning positions A2 to A12. In contrast, the L68-sfGFP insertion variant restricts this window to positions A4 to A8 (Figure 2C).

Beyond on-target window narrowing, we asked whether the L68-sfGFP insertion also mitigates Cas-independent off-target activity. Using an orthogonal R-loop assay^12^ (Figure 2D), guide RNA-independent DNA deamination was quantified within an open R-loop generated by an orthogonal *Staphylococcus aureus* Cas9 nickase (nSauCas9) targeting the GRIN2B locus, while nSpyCas9 targets CCR5. Whereas TadA8e exhibited substantial off-target editing at GRIN2B, the TadA8e-L68-sfGFP insertion variant displayed editing levels comparable to the nSauCas9 nickase-only control, indicating near-complete suppression of Cas-independent off-target deamination.

## Conclusion

This study establishes that the TadA8e scaffold can tolerate the insertion of large, autonomously folding domains at multiple sites with mild impact on overall editing efficacy. Editing behaviour depended strongly on insertion position rather than domain identity, with structurally distinct domains yielding comparable profiles at a given site (Figure 1E, F). Domain insertion thus provides a generalizable route to concurrently tune editing precision and append orthogonal functions directly within the deaminase, as shown here for a fluorescent editor with a focused editing window and reduced off-target activity (Figure 2C, D).

The broad editing window of ABE8e is attributed to the high activity and processivity of TadA8e, which allow the tethered deaminase to edit multiple adenines along the ssDNA exposed within the R-loop^4,7,10^. Consistent with this, reducing deaminase activity narrows the window and lowers off-target editing^6,8,13^. The same outcome could potentially arise from reduced expression or stability of the CRISPR base editor, either of which could result from domain insertion. While reduced deaminase activity may partially contribute to the narrowed editing window observed in this study, it alone does not account for the insertion site dependent differences we observed: domain insertion after V155 retained on-target activity at A5 comparable to the R26 and L68 variants, yet only partially reduced bystander editing and preserved a broad editing window (Figure 1E). These results point to an additional, position-specific structural mechanism underlying the editing window narrowing.

A second, alternative explanation would be that a bulky domain at a permissive site sterically constrains the spatial sampling of the deaminase over the exposed ssDNA, focusing catalysis on an optimally presented target adenine. Structural studies show that the deaminase captures its substrate by bending the ssDNA into a tRNA-like U-turn that presents a single adenine to the active site^7^. Restricting how the deaminase samples the strand would therefore limit its access to bystander adenines while preserving editing at the centrally positioned target. Importantly, the same constraint would explain the reduced guide-independent editing of the L68-sfGFP variant, as Cas-independent deamination requires the deaminase to reach transiently exposed ssDNA in trans^7,12^. Limiting its reach could therefore suppress both bystander editing and trans off-target editing. Additional experiments will be required to delineate the relative contributions of these effects to the observed narrowing of the editing window.

Unlike CRISPR-Cas effectors themselves^9,12^, which are well-amenable to optogenetic control via LOV2-mediated allostery upon domain insertion, TadA8e does not appear to be well-compatible with this control modality. This may be due to the rigid, highly thermostable fold of TadA8e^13^, which unlike CRISPR-Cas effectors, may lack intrinsic dynamics that would facilitate propagation of conformational changes throughout the domain (Supplementary Figure S2). We speculate that reducing the rigidity and thereby increasing the intrinsic dynamics of TadA8e could render it more amenable to allosteric control via receptor-domain fusion. This might be achieved, for example, by weakening fold-stabilizing interactions through mutagenesis or by applying circular permutation to the domain^14^.

Collectively, our findings demonstrate that internal domain insertion into TadA8e can simultaneously narrow the editing window, suppress Cas-independent off-target deamination, and introduce new functionality (fluorescence) - three objectives traditionally pursued through separate engineering strategies. This positions internal domain insertion as a versatile and modular approach to base-editor engineering, one that exploits the structural tolerance of TadA8e to refine editing behaviour while enabling the internal placement of orthogonal protein domains.

## Materials and Methods

### Plasmid Construction

All DNA fragments were PCR-amplified with Q5 Hot Start High-Fidelity Master Mix (NEB), all oligos were synthesized by Sigma-Aldrich. Domain inserts (*As*LOV2, PDZ, sfGFP) were introduced into the TadA8e (Addgene #138489)^4^ coding sequence immediately after the indicated residue by Golden Gate^15^ assembly (BsaI-HFv2, NEB), each flanked on both sides by a Gly-Ser-Gly (GSG) linker, of *As*LOV2 inserted after R26, Y73, and Y77, which all contain an GGSGGS linker on both sides. Spacer sequences were manually chosen or taken from previous studies^4,6,16^, gRNA plasmids were assembled by annealing complementary spacer oligonucleotides and inserting them into a SpyCas9 gRNA scaffold plasmid^17^ by Golden Gate assembly (BbsI-HF, NEB). The nSauCas9 R-loop plasmid was assembled by restriction–ligation cloning (SacI, NEB; T4 DNA ligase, Thermo Fisher). Cloning products were transformed into chemically competent TOP10 *E. coli* (Thermo Fisher), purified with the QIAprep Spin Miniprep Kit (Qiagen), and verified by Sanger sequencing (Microsynth).

### Cell Culture and Transfection

HEK293T cells were cultured in phenol red free DMEM supplemented with 10% (v/v) fetal calf serum, 1% (v/v) Penicillin-Streptomycin, and 1% (v/v) GlutaMAX at 37 °C and 5% CO_2_. Cells were passaged every 3 - 4 days to keep confluency below 90 %. For transfection, cells were seeded in 96-well plates in 100 µL with 1.25 × 10^4^ cells per well and transfected 24 h later with Lipofectamine 2000 (Thermo Fisher) according to the manufacturer’s protocol.

### Base Editing and Orthogonal R-loop Assay

For endogenous base editing, cells were co-transfected with 75 ng base-editor plasmid and 75 ng gRNA plasmid and harvested after 48 h. Cas-independent deamination was assessed with an orthogonal R-loop assay^11^. Cells were co-transfected with 66 ng base-editor plasmid, 66 ng of the corresponding base-editor gRNA plasmid (targeting *CCR5*), and 66 ng of a plasmid co-encoding n*Sau*Cas9 and a gRNA targeting *GRIN2B*. For control samples only containing *Sau*Cas9 or nSauCas9, 132 ng pBluescript II SK was added to keep the total DNA amount consistent. 72 h post transfection cells were harvested and editing efficiency analysed.

### Amplicon Sequencing and Editing Quantification

Cells were lysed in DirectPCR Lysis Reagent (VWR) with Proteinase K (∼0.2 mg/mL, Sigma-Aldrich) at 55 °C for 8 h, followed by heat inactivation at 85 °C for 45 min. Target loci were amplified directly from crude lysates with OneTaq 2× Master Mix (NEB), resolved on a 1% TAE–agarose gel, gel-purified (QIAquick Gel Extraction Kit, Qiagen), and Sanger-sequenced (Microsynth) with the forward PCR primer (Supplementary table S1). A→G editing efficiency at each protospacer adenine was quantified from the Sanger chromatograms with EditR^18^ in R (v4.3.3), counting the pam distal site of the spacer as 1. Data are reported as mean ± SD of *n* = 3 biological replicates.

### Fluorescence Microscopy

Cells were transfected with 75 ng base-editor plasmid plus 75 ng gRNA plasmid (with or without a separate episomal EGFP cassette), or with 75 ng H2B-EGFP plus 75 ng pBluescript II SK. At 48 h post-transfection, cells were imaged on a Keyence BZ-X810 microscope. GFP images were recorded with an exposure time of 0.2 s (TadA8e-L68-sfGFP and H2B-EGFP) or 0.5 s (episomal EGFP).

### Computational Inference of Insertion Sites and Data Analysis

Permissive insertion sites in TadA8e were predicted with the ProDomino algorithm^9^; additional sites were selected manually within surface-exposed loops. The ABE8e structure (PDB 6VPC)^7^ was used for site mapping and AlphaFold3^19^ for prediction of structural models of ABE:domain hybrids, both rendered in PyMOL 3.1.

## Acknowledgements

We thank all members of the Niopek and Mathony labs for helpful discussions. We thank Tobias Kreusel and Benedict Wolf for help with proDomino predictions; Sabine Aschenbrenner for help and advice on mammalian cell culture; Alexander Pattberg for cloning the initial, manually selected, *As*LOV2 insertions; and Jan Mathony for feedback on the manuscript.

## References

(1) Porto, E. M. Komor, A. C. In the Business of Base Editors: Evolution from Bench to Bedside. PLOS Biol. 2023, 21 (4), e3002071. 10.1371/journal.pbio.3002071.

(2) Gaudelli, N. M. Komor, A. C. Rees, H. A. Packer, M. S. Badran, A. H. Bryson, D. I. Liu, D. R. Programmable Base Editing of A•T to G•C in Genomic DNA without DNA Cleavage. Nature 2017, 551 (7681), 464–471. 10.1038/nature24644.

(3) Gaudelli, N. M. Lam, D. K. Rees, H. A. Solá-Esteves, N. M. Barrera, L. A. Born, D. A. Edwards, A. Gehrke, J. M. Lee, S.-J. Liquori, A. J. Murray, R. Packer, M. S. Rinaldi, C. Slaymaker, I. M. Yen, J. Young, L. E. Ciaramella, G. Directed Evolution of Adenine Base Editors with Increased Activity and Therapeutic Application. Nat. Biotechnol. 2020, 38 (7), 892–900. 10.1038/s41587-020-0491-6.

(4) Richter, M. F. Zhao, K. T. Eton, E. Lapinaite, A. Newby, G. A. Thuronyi, B. W. Wilson, C. Koblan, L. W. Zeng, J. Bauer, D. E. Doudna, J. A. Liu, D. R. Phage-Assisted Evolution of an Adenine Base Editor with Improved Cas Domain Compatibility and Activity. Nat. Biotechnol. 2020, 38 (7), 883–891. 10.1038/s41587-020-0453-z.

(5) Fu, J. Li, Q. Liu, X. Tu, T. Lv, X. Yin, X. Lv, J. Song, Z. Qu, J. Zhang, J. Li, J. Gu, F. Human Cell Based Directed Evolution of Adenine Base Editors with Improved Efficiency. Nat. Commun. 2021, 12 (1), 5897. 10.1038/s41467-021-26211-0.

(6) Chen, L. Zhang, S. Xue, N. Hong, M. Zhang, X. Zhang, D. Yang, J. Bai, S. Huang, Y. Meng, H. Wu, H. Luan, C. Zhu, B. Ru, G. Gao, H. Zhong, L. Liu, M. Liu, M. Cheng, Y. Yi, C. Wang, L. Zhao, Y. Song, G. Li, D. Engineering a Precise Adenine Base Editor with Minimal Bystander Editing. Nat. Chem. Biol. 2023, 19 (1), 101–110. 10.1038/s41589-022-01163-8.

(7) Lapinaite, A. Knott, G. J. Palumbo, C. M. Lin-Shiao, E. Richter, M. F. Zhao, K. T. Beal, P. A. Liu, D. R. Doudna, J. A. DNA Capture by a CRISPR-Cas9–Guided Adenine Base Editor. Science 2020, 369 (6503), 566–571. 10.1126/science.abb1390.

(8) Evanoff, M. Korpal, S. Krill, Z. D. Cowan, Q. T. Komor, A. C. Precise, Minimally Evolved Adenine Base Editors Generated through Mutation Reversion Analysis. Nat. Biotechnol. 2026, 1–12. 10.1038/s41587-026-03045-z.

(9) Wolf, B. Shehu, P. Brenker, L. von Bachmann, A.-L. Kroell, A.-S. Southern, N. Holderbach, S. Eigenmann, J. Aschenbrenner, S. Mathony, J. Niopek, D. Rational Engineering of Allosteric Protein Switches by in Silico Prediction of Domain Insertion Sites. Nat. Methods 2025, 22 (8), 1698–1706. 10.1038/s41592-025-02741-z.

(10) Chen, X. McAndrew, M. J. Lapinaite, A. Unlocking the Secrets of ABEs: The Molecular Mechanism behind Their Specificity. Biochem. Soc. Trans. 2023, 51 (4), 1635–1646. 10.1042/BST20221508.

(11) Doman, J. L. Raguram, A. Newby, G. A. Liu, D. R. Evaluation and Minimization of Cas9-Independent off-Target DNA Editing by Cytosine Base Editors. Nat. Biotechnol. 2020, 38 (5), 620–628. 10.1038/s41587-020-0414-6.

(12) Zhu, L. Nguyen, L. T. Bell, A. G. Krebel, T. Gillmann, K. M. Cao, Q. Oatman, H. Hariri, J. Möglich, A. Myhrvold, C. Toettcher, J. E. Multimodal Control of Cas13d Activity through Domain Insertion at an Allosteric Hotspot. Nat. Commun. 2026. 10.1038/s41467-026-73645-5.

(13) Zhu, H. Wang, L. Wang, Y. Jiang, X. Qin, Q. Song, M. Huang, Q. Directed-Evolution Mutations Enhance DNA-Binding Affinity and Protein Stability of the Adenine Base Editor ABE8e. Cell. Mol. Life Sci. 2024, 81 (1), 257. 10.1007/s00018-024-05263-7.

(14) Guo, Z. Smutok, O. Lee, G. R. Cui, Z. Qianzhu, H. Kish, M. Ergun yva, C. Wu, K. Mutschler, R. Jackson, C. J. Fiorito, M. M. Warden, A. C. Smith, O. B. Quijano-Rubio, A. Huber, T. Phillips, J. J. Otting, G. Katz, E. Baker, D. Alexandrov, K. Artificial Allosteric Protein Switches with Machine-Learning-Designed Receptors. Nat. Biotechnol. 2026, 1–11. 10.1038/s41587-026-03081-9.

(15) Engler, C. Kandzia, R. Marillonnet, S. A One Pot, One Step, Precision Cloning Method with High Throughput Capability. PLOS ONE 2008, 3 (11), e3647. 10.1371/journal.pone.0003647.

(16) Bubeck, F. Hoffmann, M. D. Harteveld, Z. Aschenbrenner, S. Bietz, A. Waldhauer, M. C. Börner, K. Fakhiri, J. Schmelas, C. Dietz, L. Grimm, D. Correia, B. E. Eils, R. Niopek, D. Engineered Anti-CRISPR Proteins for Optogenetic Control of CRISPR–Cas9. Nat. Methods 2018, 15 (11), 924–927. 10.1038/s41592-018-0178-9.

(17) Senís, E. Fatouros, C. Große, S. Wiedtke, E. Niopek, D. Mueller, A.-K. Börner, K. Grimm, D. CRISPR/Cas9-Mediated Genome Engineering: An Adeno-Associated Viral (AAV) Vector Toolbox. Biotechnol. J. 2014, 9 (11), 1402–1412. 10.1002/biot.201400046.

(18) Kluesner, M. G. Nedveck, D. A. Lahr, W. S. Garbe, J. R. Abrahante, J. E. Webber, B. R. Moriarity, B. S. EditR: A Method to Quantify Base Editing from Sanger Sequencing. CRISPR J. 2018, 1 (3), 239–250. 10.1089/crispr.2018.0014.

(19) Abramson, J. Adler, J. Dunger, J. Evans, R. Green, T. Pritzel, A. Ronneberger, O. Willmore, L. Ballard, A. J. Bambrick, J. Bodenstein, S. W. Evans, D. A. Hung, C.-C. O’Neill, M. Reiman, D. Tunyasuvunakool, K. Wu, Z. Žemgulytė, A. Arvaniti, E. Beattie, C. Bertolli, O. Bridgland, A. Cherepanov, A. Congreve, M. Cowen-Rivers, A. I. Cowie, A. Figurnov, M. Fuchs, F. B. Gladman, H. Jain, R. Khan, Y. A. Low, C. M. R. Perlin, K. Potapenko, A. Savy, P. Singh, S. Stecula, A. Thillaisundaram, A. Tong, C. Yakneen, S. Zhong, E. D. Zielinski, M. Žídek, A. Bapst, V. Kohli, P. Jaderberg, M. Hassabis, D. Jumper, J. M. Accurate Structure Prediction of Biomolecular Interactions with AlphaFold 3. Nature 2024, 630 (8016), 493–500. 10.1038/s41586-024-07487-w.

(20) Muench, P. Fiumara, M. Southern, N. Coda, D. Aschenbrenner, S. Correia, B. Gräff, J. Niopek, D. Mathony, J. A Modular Toolbox for the Optogenetic Deactivation of Transcription. Nucleic Acids Res. 2025, 53 (3), gkae1237. 10.1093/nar/gkae1237.

(21) Schindelin, J. Arganda-Carreras, I. Frise, E. Kaynig, V. Longair, M. Pietzsch, T. Preibisch, S. Rueden, C. Saalfeld, S. Schmid, B. Tinevez, J.-Y. White, D. J. Hartenstein, V. Eliceiri, K. Tomancak, P. Cardona, A. Fiji: An Open-Source Platform for Biological-Image Analysis. Nat. Methods 2012, 9 (7), 676–682. 10.1038/nmeth.2019.

