## Supplementary Information for "Domain Insertion Improves the Precision of a CRISPR Adenine Base Editor"

#### Supplementary Methods:

##### Blue-Light Illumination

For optogenetic experiments, cells were exposed to pulsed blue light in a custom-built illumination system integrated into a standard $CO_2$ incubator, as described previously[20]. Briefly, 96-well plates were illuminated from below in cycles of 5 s on and 10 s off at 5 $W/m^2$ using six high-power blue LEDs (peak emission ~460 nm) controlled by a Raspberry Pi running a custom Python script

##### Transfection

Transfection was performed as described in the main method section with following modifications. 30 ng base editor plasmid and 30 ng gRNA plasmid were cotransfected.

##### BsaHI Restriction Assay

A→G editing at position 5 of the *CCR5* protospacer  generates a BsaHI recognition site (GACGTC) (Supplementary Figure S2 A), allowing editing to be quantified by restriction digestion; editing at neighboring adenines does not alter the site. Target loci were amplified from crude lysates as described above. Per reaction, 5 µL of PCR product was combined with 2 µL of 10× rCutSmart buffer (NEB), 0.5 µL of BsaHI (NEB), and 12.5 µL of nuclease-free water (20 µL total), incubated for 1 h at 60 °C and heat-inactivated for 20 min at 80 °C. Digests were resolved on a 2% (w/v) agarose gel in 1× TBE for 40 min at 100 V, imaged, and band intensities were quantified in Fiji[21]. Editing efficiency was calculated by dividing the cleaved (edited) by the uncut (unedited) band intensities.

### Supplementary Figures:

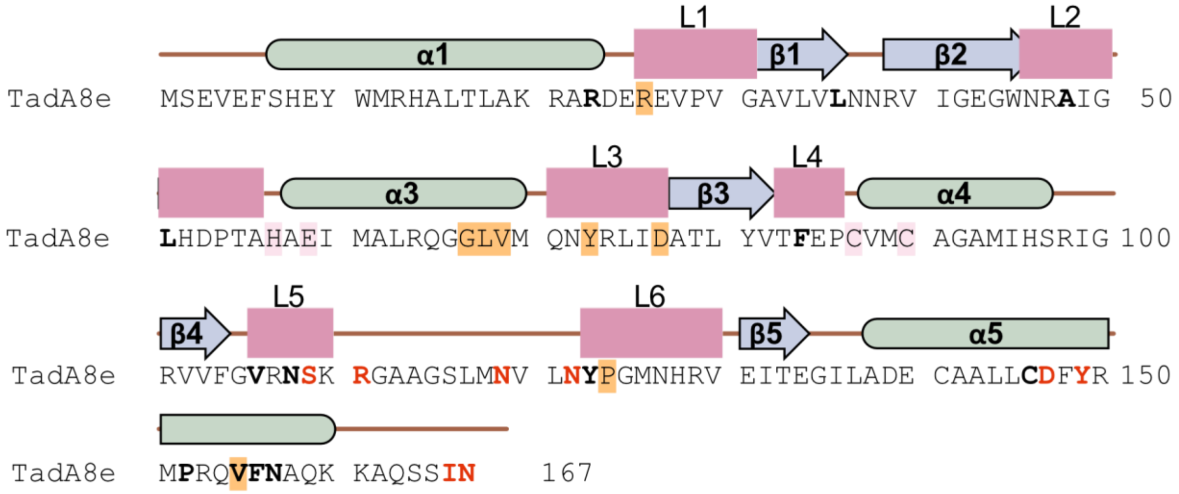


**Supplementary Figure S1. Secondary structure and engineered residues of TadA8e.** Secondary-structure elements, α-helices α1–α5 and β-strands β1–β5, are drawn above the chain, with loop regions L1-L6 shaded.Catalytic residues are boxed in pink and the eight tested insertion sites in yellow. Bold letters denote residues mutated during earlier evolution of the ABE lineage[3]; red letters denote substitutions newly acquired in TadA8e[4]. Secondary-structure assignments follow[10].

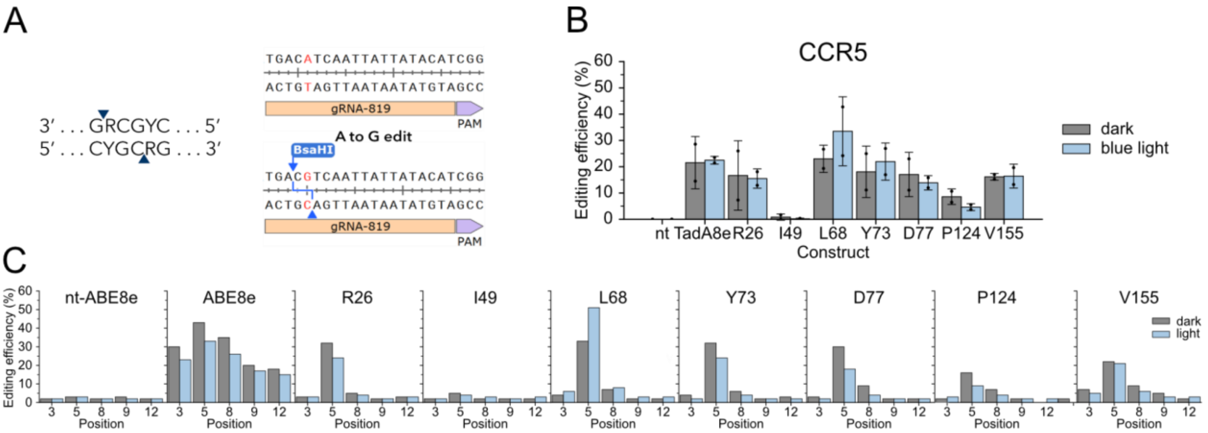


**Supplementary Figure S2. AsLOV2 insertions into TadA8e do not confer light-dependent editing.** (A) A→G editing at position 5 of the CCR5 protospacer (gRNA-819) creates a BsaHI recognition site (GACGTC), unaffected by bystander editing. (B) Position-5 editing efficiency for wild-type TadA8e and AsLOV2 insertions after the indicated residues, under dark and blue-light conditions, quantified from the relative band intensities of cleaved (edited) and uncleaved (unedited) amplicon after BsaHI digestion. Mean ± SD of n = 2 biological replicates. (C) EditR data of one of the replicates depicted in (B). Position represents the target adenine, counting the pam distal site of the spacer as 1.

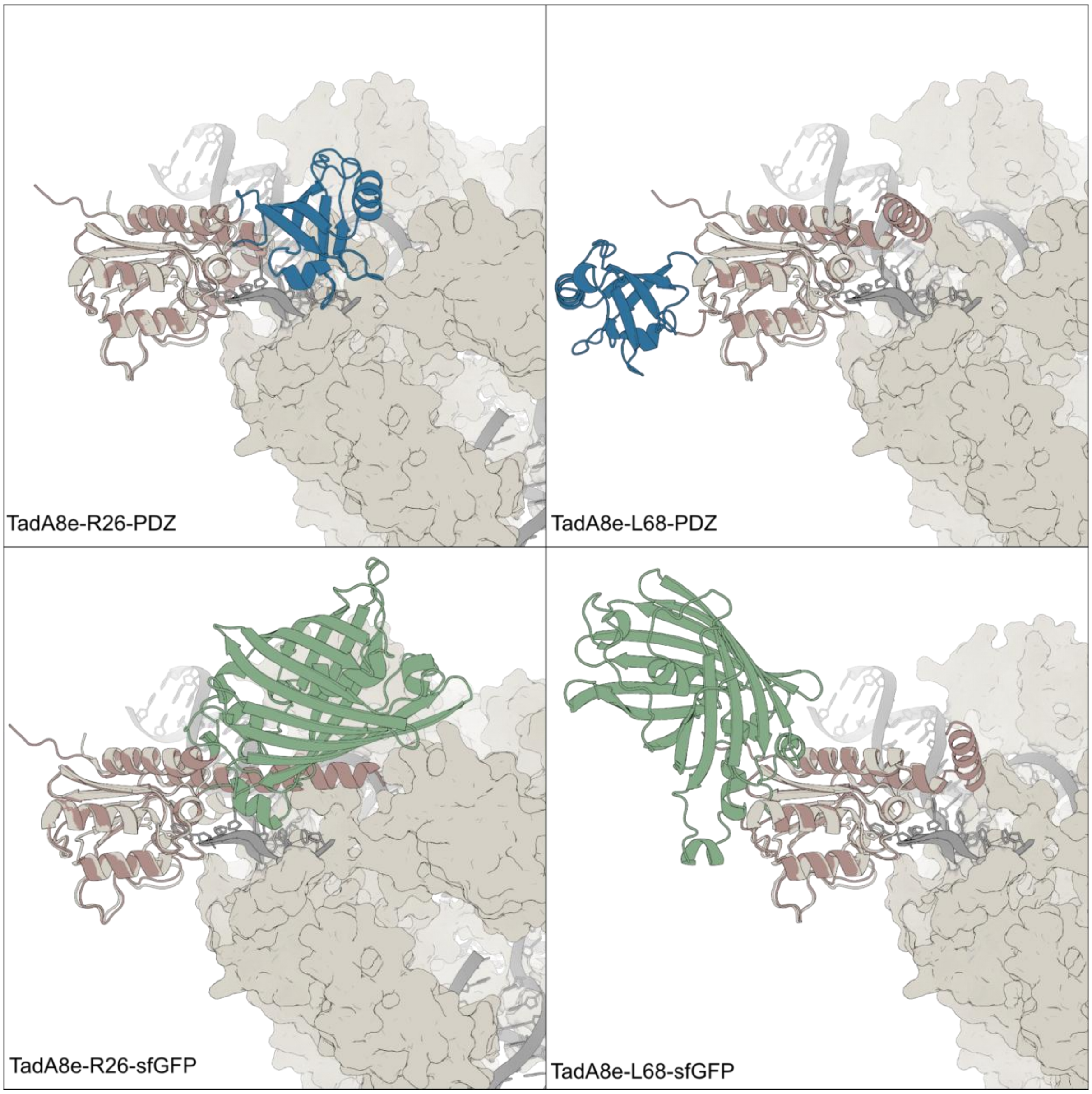


**Supplementary Figure S3. AlphaFold3 structure predictions of PDZ and sfGFP insertion variants.** Each deaminase insertion variant was predicted with AlphaFold 3[19] and superimposed on the ABE8e crystal structure by aligning the deaminase domain.The crystal structure is shown in beige with nucleic acids in grey. The predicted TadA8e domain is shown in brown, the PDZ insertion in blue, and the sfGFP insertion in green.
